# The specific AMPK activator A-769662 ameliorates pathological phenotypes following mitochondrial DNA depletion

**DOI:** 10.1101/2024.03.14.584413

**Authors:** Gustavo Carvalho, Bruno Repolês, Tran V.H. Nguyen, Josefin M.E. Forslund, Farahnaz Ranjbarian, Isabela C. Mendes, Micol Falabella, Mara Doimo, Sjoerd Wanrooij, Robert D.S. Pitceathly, Anders Hofer, Paulina H. Wanrooij

## Abstract

AMP-activated protein kinase (AMPK) is a master regulator of cellular energy homeostasis that also plays a role in preserving mitochondrial function and integrity. Upon a disturbance in the cellular energy state that increases AMP levels, AMPK activity promotes a switch from anabolic to catabolic metabolism to restore energy homeostasis. However, it is currently unclear how severe of a mitochondrial dysfunction is required to trigger AMPK activation, and whether stimulation of AMPK using specific agonists can improve the cellular phenotype following mitochondrial dysfunction. Using a cell model of mitochondrial disease characterized by progressive mitochondrial DNA (mtDNA) depletion and deteriorating mitochondrial metabolism, we show that mitochondria-associated AMPK becomes activated early in the course of the advancing mitochondrial dysfunction, before any quantifiable decrease in the ATP/(AMP+ADP) ratio or respiratory chain activity. Moreover, stimulation of AMPK activity using the specific small-molecule agonist A-769662 alleviated the mitochondrial phenotypes caused by the mtDNA depletion and restored normal mitochondrial membrane potential. Notably, the agonist treatment was able to partially restore mtDNA levels in cells with severe mtDNA depletion, while it had no impact on mtDNA levels of control cells. The beneficial impact of the agonist was also observed in cells from patients suffering from mtDNA depletion. However, the positive effects of A-769662 in the two experimental cell models appeared to involve at least partially different mechanisms. These findings improve our understanding of the effects of specific small-molecule activators of AMPK on mitochondrial and cellular function, and suggest a potential utility for these compounds in disease states involving mtDNA depletion.

## Introduction

Mitochondrial disorders are a heterogeneous group of often debilitating diseases, many of which manifest during childhood. They are caused by pathogenic variants in nuclear or mitochondrial genes that impair respiratory chain function and/or mitochondrial ATP production either directly or indirectly. The first group includes genetic variants in genes directly involved in oxidative phosphorylation (OXPHOS), while the second comprises genes whose gene products are essential for the maintenance or expression of mitochondrial DNA (mtDNA). With a few exceptions, there is currently no cure for mitochondrial diseases, and therapeutic options are thus generally limited to management of the patients’ individual symptoms (Barcelos, Emmanuele, and Hirano 2019; Pitceathly et al. 2021). Continued research into potential treatments and their mechanisms of action is thus called for.

An experimental approach that has shown promise in animal models of mitochondrial disease is the general induction of mitochondrial biogenesis, achieved for example by activating the key energy sensor and metabolic regulator, the AMP-activated protein kinase (AMPK) (Viscomi et al. 2011; Peralta et al. 2016). AMPK becomes activated upon energetic stresses manifesting as a decreased ratio of ATP to AMP and/or ADP, and phosphorylates its targets to shift metabolism from anabolism to catabolism (Steinberg and Hardie 2023). This energy-metabolic reprogramming involves upregulation of lipid and glucose breakdown, increased autophagy and mitophagy, and a boost in lysosomal and mitochondrial biogenesis (Herzig and Shaw 2018). More chronic activation of AMPK induces the expression of a core group of genes including peroxisome proliferator-activated receptor ɣ coactivator 1 α (PGC-1α), a positive regulator of mitochondrial gene expression from the nuclear genome (Malik et al. 2023; Reznick and Shulman 2006; Scarpulla 2002). In addition to transcriptional stimulation, AMPK is thought to stimulate PGC-1α activity through a combination of direct or indirect posttranslational modifications (Herzig and Shaw 2018), consequently promoting the expression of mitochondrial transcription factor A (TFAM), the high-mobility group protein that is required for the maintenance and expression of mtDNA (Larsson et al. 1998; Parisi and Clayton 1991; Garstka et al. 2003). Increased TFAM levels lead to elevated mtDNA copy number, allowing for the overall expansion of functional mitochondria (Ekstrand et al. 2004; Bonekamp et al. 2021). Thus, activation of the AMPK-PGC-1α axis increases mitochondrial biogenesis to augment respiratory capacity in response to a low cellular energy state.

In line with this concept, the pharmacological activation of AMPK using the agonist 5-aminoimidazole-4-carboxamide ribonucleoside (AICAR) has been shown to improve muscle function in different models of mitochondrial myopathy caused by complex IV (CIV) defects (Peralta et al. 2016; Viscomi et al. 2011). However, neither of the above-cited studies found evidence for the expected increase in mitochondrial biogenesis in terms of elevated mtDNA copy number or mitochondrial mass, leaving the mechanisms underlying these positive effects of AICAR somewhat unclear (Viscomi et al. 2011; Peralta et al. 2016). *In vivo*, AICAR is converted to the inosine pathway intermediate ZMP that acts as an AMP analog. In addition to binding and stimulating AMPK, ZMP modulates the activity of other AMP-regulated enzymes such as glycogen phosphorylase and fructose-1,6*-*bisphosphatase (Longnus et al. 2003; Young, Radda, and Leighton 1996; Vincent et al. 1991). Therefore, some of the positive effects of AICAR in models of mitochondrial disease might potentially be attributable to AMPK-independent actions of the drug.

More specific, non-AMP-mimetic AMPK agonists such as A-769662 and compound 991 have been developed (Cool et al. 2006; Xiao et al. 2013; Göransson et al. 2007), but have not yet been as widely adopted as AICAR. Therefore, their impact on mitochondrial function and especially mtDNA levels has not been conclusively determined. However, A-769662 and other specific AMPK agonists showed positive effects in a panel of patient-derived cell lines with genetically heterogeneous forms of mitochondrial defects (Moore et al. 2020). The group of cell lines benefiting from AMPK agonists included not only cell lines with direct defects in the OXPHOS machinery, but also ones manifesting mitochondrial dysfunction due to defects in mtDNA maintenance. Moreover, a recent study reported an increase in both mitochondrial mass and mtDNA copy number in HEK293 cells treated with compound 991 (Malik et al. 2023). These findings raise the possibility that specific AMPK agonists may be beneficial even in cases with indirect OXPHOS defects resulting from decreased mtDNA copy number, *i.e.* mtDNA depletion.

MtDNA depletion syndromes (MDS) are associated with severe, infantile-onset mitochondrial dysfunction with highly tissue-specific manifestations typically caused by pathogenic variants of enzymes involved in nucleotide metabolism or mtDNA replication, such as the mtDNA polymerase ɣ (Polɣ) (Suomalainen and Battersby 2018; Carvalho et al. 2021; El-Hattab and Scaglia 2013). The Polɣ holoenzyme consists of one catalytic PolɣA subunit that associates with a homodimer of accessory PolɣB subunits which improve processivity of the holoenzyme (Yakubovskaya et al. 2006; Johnson et al. 2000; Carrodeguas et al. 2001; Fan et al. 2006). The catalytic activity of PolɣA relies on the evolutionarily conserved aspartate residues D890 and D1135 (Spelbrink et al. 2000). Accordingly, inducible overexpression of the PolɣA^D890N^ or PolɣA^D1135A^ variants in cultured human cells containing an endogenous wildtype copy of the *POLG* gene exerts a dominant-negative phenotype with prominent mtDNA replication stalling and subsequent rapid mtDNA depletion, acting as an excellent cell model for MDS progression (Wanrooij et al. 2007; Jazayeri et al. 2003).

In this study, we use a Flp-in T-Rex 293 cell line with inducible expression of PolɣA^D1135A^ to address the role of AMPK activity during escalating mitochondrial dysfunction caused by severe mtDNA depletion. We first delineate the timing of events during the advancing decline in mtDNA levels caused by the expression of the dominant-negative D1135A variant. We demonstrate that AMPK activation is an early event that occurs before the cellular energy state or mitochondrial membrane potential is measurably affected, confirming that AMPK successfully restores energy homeostasis during the early stages of mitochondrial dysfunction. Notably, the observed activation is limited to AMPK molecules in the mitochondrial fraction of the cell, while cytosolic AMPK is not affected. Next, we show that AMPK contributes to the maintenance of mitochondrial membrane potential even under basal conditions, and that stimulation of AMPK activity using the specific agonist A-769662 is sufficient to fully restore the mitochondrial membrane potential in mtDNA-depleted cells. This positive impact of A-769662 can be at least partially attributed to a small increase in the levels of mtDNA and respiratory chain subunits; however, these benefits were insufficient to overcome the proliferation defect seen in conjunction with the mtDNA depletion. Treatment with the AMPK agonist also elevated mitochondrial membrane potential in cell lines derived from patients suffering from Polɣ-associated mitochondrial myopathy, but then through an mtDNA-independent mechanism. Our findings also uncover differential effects of the AMPK agonist in control cells and ones suffering from severe mtDNA depletion.

## Materials and Methods Cell culture

Inducible Flp-in^TM^ T-Rex^TM^ 293 cell lines (female) carrying one integrated copy of *myc*-*POLG* (wt or D1135A) were generated previously (Wanrooij et al. 2007). Cells were grown in low-glucose DMEM medium (Gibco) containing 1 g/L glucose, 4 mM GlutaMAX^TM^, 1 mM sodium pyruvate, 50 µg/mL uridine, 10% heat-inactivated fetal bovine serum (Gibco) and were cultivated in a 37°C humidity incubator at 8% CO_2_. Expression of myc-Polɣ was induced by the addition of 3 ng/mL doxycycline (Sigma) to the medium for the time periods indicated in the legends. Doxycycline-containing media was exchanged every 2-3 days. When indicated in the figures, AMPK activity was modulated by the administration of its pharmacological agonist, A-769662 (Sigma), while control, “untreated” cells received an equivalent volume of DMSO. The ρ^0^ cells used as a control were generated by 44-day treatment of the *myc*-*POLG^D1135A^* Flp-in^TM^ T-Rex^TM^ 293 cell line with 150 ng/ml ethidium bromide, and the lack of mtDNA confirmed by qPCR (Fig. S1C). Growth of the ρ^0^ cells was in high-glucose (4.5 g/L) DMEM media supplemented as above.

Human primary fibroblasts derived from two patient biopsies bearing the PolɣA mutations Y955C or V1106A, both homozygous, were cultivated in the same conditions as the Flp-in T-Rex 293 cells. The control cells used in parallel were sex- and age-matched. The research presented in this article complies with all relevant ethical regulations. Written informed consent was obtained from all participants or their guardians. This study was approved by the Queen Square Research Ethics Committee, London, UK (09/H0716/76).

## Cell proliferation

Growth curves were generated by seeding 20 000 cells/well in 12-well plates. Each day, cells from three wells per condition were trypsinized, cells were resuspended in medium, and counted using an automated cell counter (Countess II FL, Invitrogen). Trypan blue was used to assess cell viability and to exclude dead cells during cell counting.

## MtDNA copy number

Total DNA was isolated from cells at 80% confluency in a 6-well plate using the NucleoSpin® Tissue DNA isolation kit (Macherey-Nagel), according to the manufacturer’s instructions. MtDNA copy number was analyzed essentially as previously described (Repolês et al. 2021). Short targets in both the mtDNA and the nuclear DNA were quantified in duplicate by quantitative real-time PCR using 6-12 ng total DNA in a 20 μl reaction containing 0.2 μM forward and reverse primers and 10 μl of 2 × SyGreen Mix (qPCRBIO #PB20.14-05) in a LightCycler 96 instrument (Roche). Primer pairs targeted the 16S rDNA region in the mtDNA (forward GTCAACCCAACACAGGCATGCT, reverse CGTGGAGCCATTCATACAGGTCC); and a region of the single-copy XPC gene in the nuclear genome (forward GCTGGACCATCTGCTGAACCC, reverse TCCTTCCACCCCTCACCTTATGT). All primers were confirmed to target only the single target region of the genome using the BLAST tool. Cycling conditions were: 95°C 180 sec, 35 cycles of (95°C 10 sec, 57°C 10 sec, 72°C 20 sec with signal acquisition), melting curve (95°C 5 sec, 65°C 60 sec, heating to 97°C at 0.2°C/sec with continuous signal acquisition). C_q_ values determined by the LightCycler 96 software (Roche) were used to calculate the copy number of mtDNA relative to nuclear DNA using the Pfaffl method (Pfaffl 2001) and plotted with GraphPad Prism software (version 10; GraphPad Software Inc, CA).

## Protein extraction and Western blotting

Cells were grown to 80% confluence in 6-well plates, resuspended in cold phosphate buffered saline (PBS) and pellets were flash-frozen in liquid nitrogen and kept in -80°C freezer. Total protein extracts were prepared by resuspending the cell pellets in RIPA buffer (50 mM Tris-HCl pH 8.0, 150 mM NaCl, 1% NP-40, 0.5% Na-deoxycholate, 0.1% SDS) containing protease and phosphatase inhibitors (100 µM AEBSF-HCl, 4 µM aprotinin, 70 µM E-64, 110 µM Leupeptin, 5 µM pepstatin A, 5 mM NaF, 1 mM Na_3_VO_4_). DNA was sheared by passing the samples 5 times through a 27G needle. Samples were left 30 min on ice and then centrifuged for 10 min at 17 000 × g at 4°C. Supernatants containing cell lysates were transferred to new tubes and protein concentrations were measured using the BCA method (PierceTM BCA Protein Assay kit, Thermo Scientific). Total protein extracts (10-20 µg) were diluted in Laemmli buffer and boiled at 95°C for 5 min. Samples were loaded onto 4-20% MiniPROTEAN® TGX^TM^ stain-free polyacrylamide gel (Bio-Rad) and resolved by SDS-PAGE, containing Tris-Glycine-SDS buffer (50mM Tris, 250mM glycine, 0.08% SDS). Samples used for visualization of the OXPHOS complexes using the OXPHOS antibody cocktail were loaded onto 4-12% NuPAGE^TM^ Bis-Tris (ThermoFischer) and 1×MOPS was used as running buffer. Proteins were then transferred to a PVDF membrane (Amershan Hybond) in presence of Tris-Glycine (no SDS) buffer containing 20% ethanol. The membranes were blocked with Tris-buffered saline containing 0.1% Tween-20 (TBS-T) and 5% skimmed milk for 1h at room temperature (RT) and incubated overnight at 4°C with primary antibodies diluted in 5% milk TBS-T. Primary antibodies used in this study are: p-AMPKα Thr172 (1:1.000, Cell signaling #2535), AMPKα (1:500, Cell signaling #2603), p-ACC (1:1000, Cell signaling #3661), ACC (1:1000, Cell signaling #3662), p-ATM (1:1000, Santa Cruz sc-47739), ATM (1:1000, Santa Cruz sc-377293), OXPHOS cocktail (1:2000, Invitrogen 45-8199), myc-tag (1:1000, Invitrogen MA1-21316), GAPDH (1:10000, Invitrogen MA5-15738), VDAC/Porin (1:5000, Abcam ab14734), COX2 (1:10000, Invitrogen A6404), TFAM (1:10000, Abcam ab176558), β-actin (1:15000, Sigma A5441). Secondary antibodies (anti-mouse (1:10000-20000, Pierce #31430) and anti-rabbit (1:20000, Themo Scientific #31460)) used were HRP-linked diluted in 5% milk TBS-T, followed by 1 h incubation at RT with the membrane. After extensive wash with TBS-T, the blots were developed by chemiluminescence (ECL Bright or ECL SuperBright, Agrisera) and images were taken using ChemiDoc Imaging System (Bio-Rad). Image analysis and quantifications were done using ImageJ software (version 1.54g).

## Cell fractionation

To determine the sub-cellular localization of (p)AMPK, two 10 cm plates per condition were grown to 80% confluence, harvested, washed twice with cold PBS, and then resuspended and incubated in 0.1 × hypotonic buffer (4 mM Tris-HCl pH 7.8, 2.5 mM NaCl, 0.5 mM MgCl_2_ containing protease and phosphatase inhibitors) for 10 min at 4°C. Cells were homogenised in a 1 mL glass/glass Wheaton tight-fitting dounce (Active Motif) until ca 90% of cells were disrupted, and samples made isotonic by addition of 1/10 vol of 10 × homogenization buffer. Nuclear, cytosolic, and crude mitochondrial fractions were obtained after differential centrifugation. Briefly, nuclear fractions were pelleted by centrifugation at 1 200 × g for 3 min, and the supernatant containing cytosolic and mitochondrial fractions was transferred to new tubes. The centrifugation was repeated twice to remove any remaining nuclear material. From the resulting supernatant, mitochondria were pelleted by centrifugation at 17 000 × g for 10 min and the supernatant was collected as the cytosolic fraction. Proteins from the different fractions were extracted and quantified as described above.

## Blue native PAGE (BN-PAGE) and CIV in-gel activity

To assess the integrity and function of the mitochondrial complexes and supercomplexes, mitochondria were isolated from harvested cells, solubilized, and resolved in blue native PAGE to further perform CIV in-gel activity determine and protein levels by immunoblotting according to a protocol adapted from (Frezza, Cipolat, and Scorrano 2007). Briefly, cells from two 150 mm plates per condition were harvested at 90% confluence, suspended in IBc buffer (10 mM Tris-MOPS, 1 mM EGTA, 0.2 M sucrose) containing protease and phosphatase inhibitors, and homogenized in a 15 mL glass/glass Wheaton tight-fitting dounce (Active Motif) until they were 55-65% disrupted (positive for Trypan blue staining according to automated cell counting (Countess II FL, Invitrogen). Mitochondrial pellets were obtained by differential centrifugation, resuspended in IBc buffer and the proteins quantified using the BCA method. Aliquots of mitochondrial fraction were flash-frozen in liquid nitrogen and stored at -80°C for up to 2 weeks. On the day of the assay, mitochondrial pellets were thawed on ice and solubilized in NativePAGE^TM^ buffer (Invitrogen) containing 4% digitonin and protease inhibitors. After 1.5 h incubation with solubilization buffer, the samples were centrifuged at 16000 × g for 20 min and the supernatant containing solubilized mitochondria was mixed with NativePAGE^TM^ G-250 sample additive (Invitrogen). Mitochondrial samples (100 µg for in-gel activity; 25 µg for WB) were loaded onto NativePAGE^TM^ 3-12% Bis-Tris gel (Invitrogen) and resolved by BN-PAGE containing NativePAGE^TM^ Running buffer (Invitrogen) and NativePAGE^TM^ cathode buffer additive (Invitrogen). NativeMark™ Unstained Protein Standard (Invitrogen) was used for molecular weight estimation of protein in native gel electrophoresis. To assess the activity of CIV, the gel was incubated overnight with substrate solution (50 mM Na-phosphate buffer pH7.4, 0.5 g/L 3,3’-diamidobenzidine tetrahydrochloride, 1.0 g/L bovine cytochrome *c*, 2 mg/L catalase, 75 g/L sucrose) until the brown signal appeared on the gel. Images of the gel were recorded using a high-quality imager (Epson Perfection V700 Photo).

## Mitochondrial membrane potential (MMP)

Cells were grown to 70-80% confluence in 6-well plates. For each condition, one well was used for tetramethylrhodamine ethyl ester (TMRE) staining and another for 10-N-nonyl acridine orange (NAO) staining. To assess MMP, adherent cells were incubated with medium containing 200 nM TMRE for 30 min at 37°C. After incubation, cells were trypsinized and resuspended in PBS containing 1% BSA, and TMRE fluorescence was determined by flow cytometry (BD Accuri^TM^ C6 Plus, BD Biosciences). TMRE fluorescence was also measured in cells further treated with the mitochondrial uncoupler BAM-15 (60 µM) for 30 min at 37°C to exclude non-mitochondrial TMRE fluorescence. Mitochondrial mass was estimated by the fluorescence of cells stained with 10 nM NAO for 30 min at 37°C. The MMP was calculated by subtracting the TMRE fluorescence of BAM-15 uncoupled cells from the respective coupled ones (ΔTMRE = TMRE_coupled_ – TMRE_uncoupled_) and the result was normalized by the NAO fluorescence (ΔTMRE/NAO).

## AMPK silencing

AMPK silencing was achieved via cell transfection with AMPKα1/α2 siRNA (#45312, Santa Cruz), and scrambled siRNA (#4390844, ThermoFisher) was used as negative control. Shortly, 50% confluent wells (6-well plate) were transfected with 100 pmol siRNA and Lipofectamine® RNAiMAX (Invitrogen) prepared in OptiMEM® (Gibco) medium, according to manufacturer. After 24 h transfection, the medium was replaced with fresh medium containing either DMSO or 100 µM A-769662, and cells were harvested 48 h later for MMP measurement and WB analysis.

## Nucleotide pool measurement

Nucleotide (dNTP and NTP) pools were measured in a single run using a recently developed isocratic reverse phase HPLC-based technique (Ranjbarian et al. 2022), with minor modifications for the analysis of the relative levels of AMP, ADP and ATP (Purhonen, Hofer, and Kallijärvi 2024; Debar et al. 2023). Briefly, cells were cultivated in 10 cm plates to 50-75% confluence, washed with ice-cold PBS, and scraped off the plates in the presence of 0.5 mL ice-cold 80% (v/v) methanol in water. Cells were centrifuged for 1 min at 17 000 × g in a cooled centrifuge, and the supernatants containing the free nucleotides were collected in new tubes and stored at -80°C. The cell extracts were purified by solid phase extraction (SPE) using Oasis WAX 3cc cartridges (Waters Corporation, Milford, MA, USA) in a similar manner as described before for the measurement of NTPs, dNTPs, and ADP (Ranjbarian et al. 2022), but with some minor modifications in the SPE step in experiments where also AMP was measured (Debar et al. 2023). The SPE-purified samples were subsequently analyzed by HPLC run at 1 ml/min using a 150 mm × 4.6 mm SunShell C18-WP HPLC column from ChromaNik Technologies Inc (Osaka, Japan). The aqueous mobile phase contained 5.8% (v/v) acetonitrile, 0.7 g/L tetrabutylammonium bromide as ion pairing agent and varying concentrations of potassium phosphate at pH 5.6. The analysis of NTPs and dNTPs was performed with the isocratic Fast Protocol (Ranjbarian et al., 2022), whereas the analysis of AMP, ADP and ATP was performed with a phosphate gradient (Purhonen, Hofer, and Kallijärvi 2024).

## Cell cycle

Cells were grown to 70-80% confluence in 6-well plates, harvested and fixed with ice-cold 70% ethanol, and kept in freezer -20°C up to 1 week. For DNA staining, cells were washed with PBS and resuspended in staining solution containing 0.02 mg/mL propidium iodide, 0.1% Triton X-100, and 0.2 mg/mL RNAse in PBS. Cells were incubated with staining solution for 30 min at RT. Cell cycle was assessed by flow cytometry (BD Accuri^TM^ C6 Plus, BD Biosciences) and data was analysed using FlowJo^TM^ BD software.

## Cell death assay

Levels of cell death were measured in cells expressing Polγ^WT^ and Polγ^D1135A^ for 6 days using the FITC/Annexin V Dead Cell Apoptosis kit (Molecular Probes, Invitrogen) according to the manufacturer’s instructions. Uninduced Polγ^WT^ cells treated with 10 μM of camptothecin for 16 hours were used as a positive control for cell death. Cells were harvested, pelleted by centrifugation, washed once with 1 × PBS and resuspended in PBS to a concentration of 10^6^ cells/ml. For each measurement, 10^5^ cells were combined with 5 μl FITC Annexin V solution and 1 μl of 100 μg/mL propidium iodide solution. The samples were incubated at RT, protected from light, for 15 min. After incubation, 400 μL of annexin binding buffer was added to the mixture, gently mixed, and kept on ice. Samples were measured immediately after the addition of the buffer on a BD Accuri C6 Plus (BD Biosciences) flow cytometer. Unstained and uninduced cells were used as a negative control.

## Graphs and statistical analysis

All graphs and statistical analyses were generated with GraphPad Prism software (version 10; Graphpad Software Inc, CA). For experiments comparing uninduced and induced Flp-in T-REx 293 cells across multiple timepoints, data of all induced samples was normalized to the induced sample at day 0 to eliminate technical variation arising from different doxycycline preparations. To clearly indicate this, a vertical dotted line is used to divide the graphs into two halves, where data sets in each half are normalized to their respective 0-day sample. Results are expressed as mean ± standard deviation, and statistical differences (unpaired t-test) indicated by asterisks (* P≤0.05, ** P≤0.01, *** P≤0.001, **** P≤0.0001). The number of biological replicates (n) is indicated in the figure legends.

## Results

### The progressive mtDNA depletion induced by Polɣ^D1135A^ expression impairs OXPHOS and cell proliferation within 4 to 6 days

We used the previously established Flp-In T-REx 293 cell line to express the dominant-negative D1135A variant of the mitochondrial DNA polymerase PolɣA (henceforth referred to as Polɣ^D1135A^) in a doxycycline-inducible manner (Wanrooij et al. 2007). A similar cell line expressing the wildtype PolɣA (Polɣ^wt^) was used as a control. To better mimic physiological energy metabolism and mitochondrial function, the cells were maintained in low-glucose media (5 mM glucose). The expression of Polɣ^wt^ and Polɣ^D1135A^ was induced by the addition of 3 ng/ml of doxycycline, a low concentration that was selected because it decreased mtDNA levels to below 15% of starting levels by day 3 in Polɣ^D1135A^-expressing cells (Fig. 1A). No leaky expression of Polɣ^wt^ or Polɣ^D1135A^ was observed in the absence of doxycycline (Fig. S1A-B).

**Figure 1.**
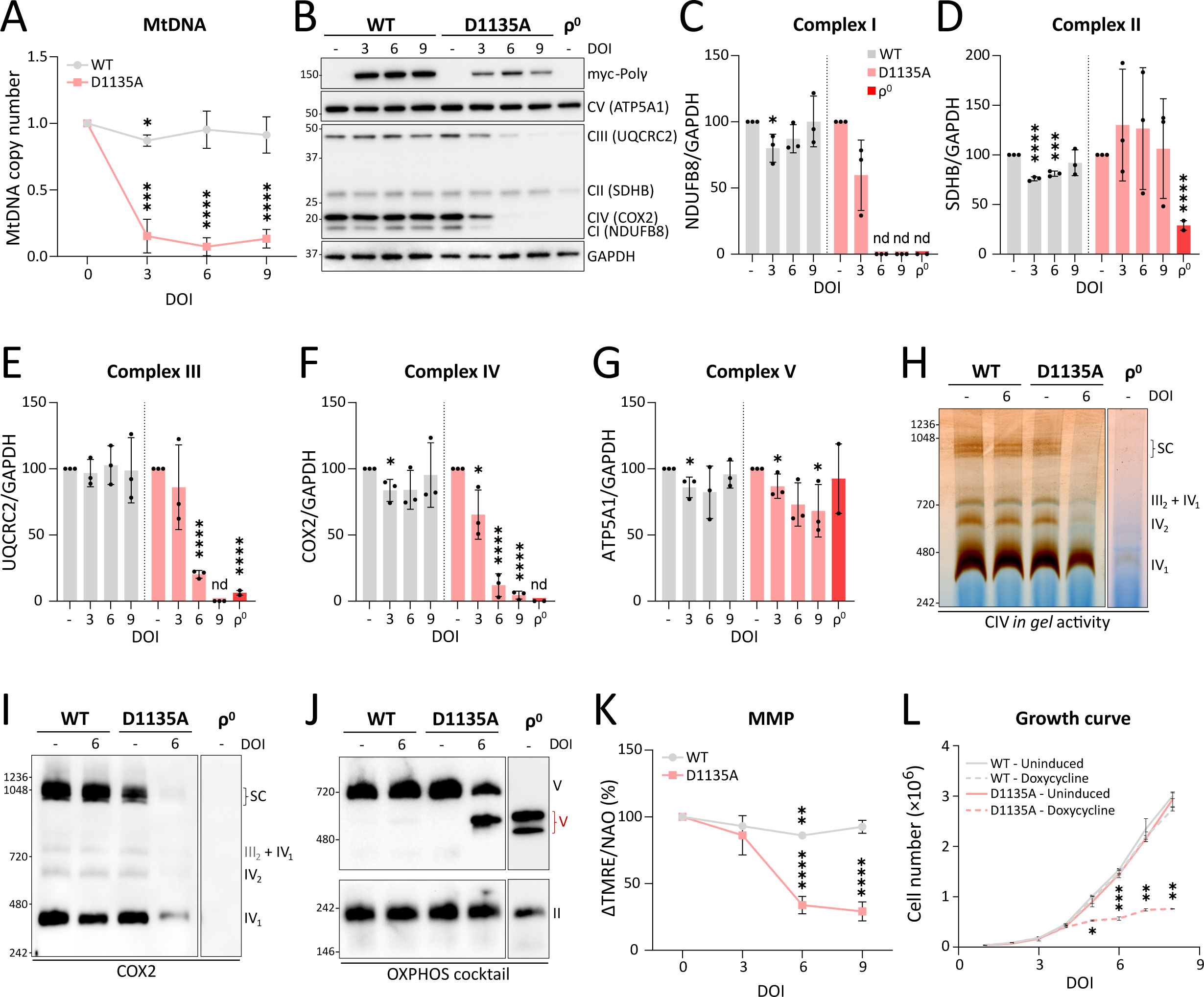
Progressive loss of mitochondrial function in a model of mtDNA depletion. **(A)** Relative mtDNA copy number following the overexpression of Polγ^wt^ (*grey*) and the dominant-negative Polγ^D1135A^ (*pink*) over the course of 3, 6 and 9 <u>d</u>ays <u>o</u>f induction (DOI) with 3 ng/mL doxycycline. The relative mtDNA copy number was normalised to the respective uninduced cells (0 DOI); n=3. **(B)** Expression of subunits of the mitochondrial respiratory complexes and the ATP synthase (complexes I to V) after 3, 6, and 9 days of induction of Polγ^wt^ and Polγ^D1135A^ (indicated as myc-Polγ) and HEK293 ρ^0^ cells were assessed by WB using an OXPHOS cocktail antibody. The position of protein molecular weight markers (kDa) is indicated on the left and protein names (and their respective respiratory complexes) are indicated on the right. A representative image of n=3. **(C, D, E, F and G)** Quantification of the blots shown in **(B)**, with Polγ^wt^ (*grey*), Polγ^D1135A^ (*pink*), and ρ^0^ cells (*red*). Protein levels were normalized against GAPDH. **(H)** The function of CIV and supercomplexes containing CIV was assessed by *in-gel* activity assay after BN-PAGE from the indicated cell lines. Representative image of n=3. **(I)** Immunoblotting with COX2 antibody to assess the integrity of CIV following BN-PAGE. Representative image of n=3. **(J)** Immunoblotting with OXPHOS cocktail antibody to detect CI, CII and CV in a BN-PAGE gel. The following complexes and supercomplexes are indicated: (IV_1_) CIV monomer; (IV_2_) CIV dimer; (III_2_ + IV_1_) CIII dimer and CIV monomer; (SC) respiratory <u>s</u>uper<u>c</u>omplexes formed by CI, CIII and CIV; (V) complex V. In cells expressing Polγ^D1135A^ for 6 days and in ρ^0^ cells the accumulation of partially assembled CV (in red) is detected. Representative image of n=3. **(K)** Mitochondrial membrane potential (MMP) was estimated by flow cytometry using TMRE dye. Non-mitochondrial TMRE fluorescence was subtracted from cells treated with the mitochondrial uncoupler BAM15 (ΔTMRE). TMRE signal was adjusted to mitochondrial mass estimated by staining mitochondria with 10-N-nonyl acridine orange (NAO). Relative MMP is expressed as ΔTMRE/NAO; n=3. **(L)** Growth curve of uninduced cells or induced cells overexpressing either Polγ^wt^ or Polγ^D1135A^ (*doxycycline*) was obtained by counting the absolute number of cells over the course of 8 days; n=2. (n, number of biological replicates; nd, not determined; DOI, days of induction; * P≤0.05; ** P≤0.01; *** P≤0.001; **** P≤0.0001).

To follow the dynamics of the mitochondrial and cellular effects of mtDNA depletion, the expression of Polɣ^D1135A^ was induced, and samples were collected 3, 6 and 9 days after induction. Cells expressing Polɣ^D1135A^ exhibited a considerably decreased copy number of 8-15% of starting levels across all three post-induction timepoints (Fig. 1A), while the protein levels of respiratory chain (RC) subunits showed a decreasing trend over the time course (Fig. 1B-G). Subunits of complex IV and V were mildly affected already on day 3, and by day 6, all complexes except for the fully nuclear-encoded CII showed diminished protein levels. In line with the trend in the Flp-In T-REx 293 ρ^0^ control cells, subunits of complexes I, III and IV were most severely affected, while the decrease in ATP5 (CV) levels was more moderate (ca 70% of initial protein levels remaining on day 9; see Fig. S1C-D for validation of the ρ^0^ cells). In contrast to Polɣ^D1135A^, the expression of Polɣ^wt^ had little or no impact on copy number or RC complex levels, confirming that the observed effects were not related to an overwhelmed protein import system upon overexpression of a mitochondrially-targeted protein. Blue-native analysis confirmed a pronounced loss of assembled RC complexes and supercomplexes in Polɣ^D1135A^ cells on day 6 of induction, including the appearance of a smaller subcomplex of CV that was also observed in ρ^0^ cells by us and others (Fig. 1H-J)(Herrero-Martin et al. 2008). CIV activity on day 6 of induction was notably decreased compared to uninduced Polɣ^D1135A^ cells (Fig. 1H-J). Accordingly, the mitochondrial membrane potential (MMP), measured using the cationic dye tetramethylrhodamine methyl ester perchlorate (TMRE) and normalized to mitochondrial mass assessed by 10-N-nonyl acridine orange (NAO), diminished to 35 % of starting levels by day 6 (Fig. 1K). Cell proliferation stagnated already on day 5, suggesting that the decrease in OXPHOS and MMP observed in day 6 samples may be physiologically relevant already on day 5 when no samples were collected for protein or MMP analysis (Fig. 1L).

### Mitochondria-associated AMPK is activated early on during the course of the mitochondrial dysfunction

To further assess the effects of mtDNA depletion on energy metabolism, we next used HPLC to quantify the levels of adenine nucleotides in Polɣ^D1135A^ or Polɣ^wt^-expressing cells before induction, as well as 3 or 6 days after the addition of doxycycline. In line with the observed MMP defect, the ADP level in Polɣ^D1135A^-expressing cells was elevated at day 6, and AMP levels showed a reproducible increase that however failed to reach statistical significance (p=0.117; Fig. 2A). The cells’ energy status, appraised by the ratio of ATP to the sum of AMP and ADP, thus decreased 2.5-fold compared to uninduced Polɣ^D1135A^ cells (computed ATP/(AMP+ADP) ratio of 12.0 and 4.7 in uninduced and day-6 cells, respectively). Further analysis of nucleotide levels revealed a decline in all pyrimidine nucleoside triphosphates (CTP, UTP, dCTP and dTTP) in Polɣ^D1135A^ cells, which is an expected consequence of the respiratory chain deficiency impairing pyrimidine synthesis (Fig. S1E-F) (King and Attardi 1989; Grégoire et al. 1984). The Polɣ^D1135A^ cells also exhibited activation of the ATM checkpoint kinase and an altered cell cycle profile (Fig. S1G-I). No signs of increased apoptosis were detected (Fig. S1J). These findings are in good agreement with the previously-reported cell cycle alterations and ATM activation observed in response to mtDNA instability, and the fact that mitochondrial dysfunction can lead to nuclear DNA instability through a host of different mechanisms (Veatch et al. 2009; Hämäläinen et al. 2019; Marcon et al. 2022; Cao et al. 2022).

**Figure 2.**
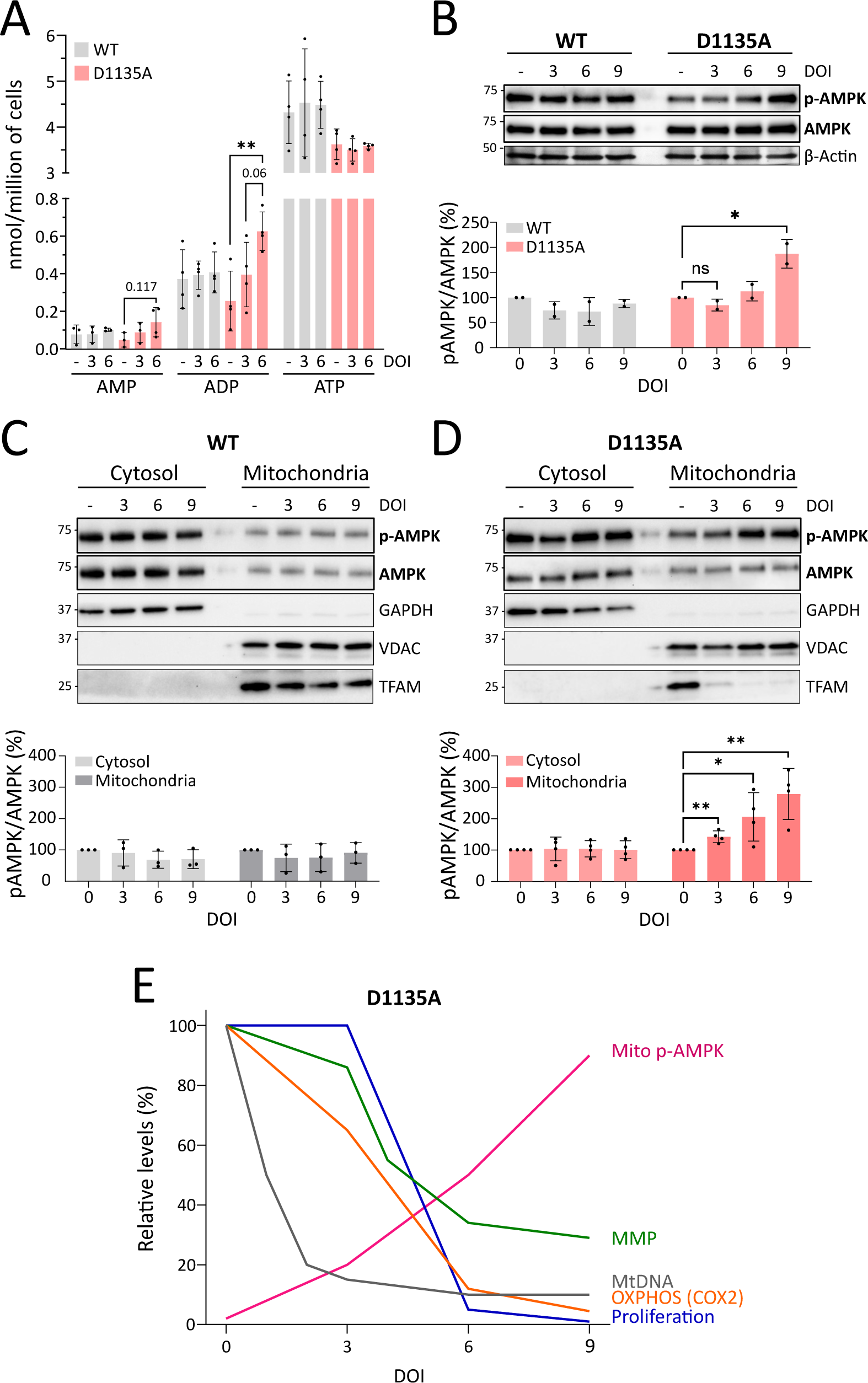
MtDNA loss leads to energy imbalance and activation of mitochondria-associated AMPK. **(A)** Levels of AMP, ADP and ATP in cells either uninduced (day 0) or expressing PolG^wt^ (*grey*) or PolG^D1135A^ (*pink*) for 3 and 6 days; n=4. **(B)** Immunoblotting indicates levels of phosphorylated AMPK (p-AMPK) and total AMPK in cells overexpressing PolG^wt^ and PolG^D1135A^. β-Actin was used as loading control. A representative image of n=2 is shown. **(C, D)** Cell fractionation was performed to show the subcellular localization of AMPK. Immunoblotting shows the distribution of p-AMPK and AMPK in the cytosolic and mitochondrial fraction of cells overexpressing PolG^wt^ (n=3) and PolG^D1135A^ (n=4). Quantification of the p-AMPK/AMPK ratios is shown in the bottom panel. GAPDH and VDAC are shown as a purity control for the cytosolic and mitochondrial fractions, respectively. TFAM levels provide an indirect estimation of mtDNA copy number (see also Fig. 1A). **(E)** A schematic summarizing the advance of the cellular phenotypes associated with the progressive loss of mtDNA shown in figures 1 and 2. (n, number of biological replicates; ns, not significant; DOI, days of induction; * P≤0.05; ** P≤0.01).

Given the altered ATP/(AMP+ADP) ratio in Polɣ^D1135A^-expressing cells, we next examined the status of the AMPK energy sensor, a fraction of which localizes to the mitochondrial outer membrane (Drake et al. 2021). Specifically, we were interested in understanding when over the course of the progressive mitochondrial dysfunction in the Polɣ^D1135A^ cells AMPK becomes activated and thus phosphorylated. Analysis of AMPK phosphorylation on threonine-172 in whole-cell lysates suggested that AMPK was not activated until day 9 of induction (Fig. 2B). However, evaluation of AMPK in different subcellular localizations revealed that mitochondria-associated AMPK underwent activatory phosphorylation already on day 3, while the phosphorylation state of cytosolic AMPK remained at basal levels throughout the time course (Fig. 2C-D). Notably, the energy stress in Polɣ^D1135A^ cells did not alter the partitioning of AMPK between the cytosolic and crude mitochondrial fraction, indicating no significant recruitment of additional AMPK complexes to the mitochondria (Fig. 2D). As expected, the induced expression of Polɣ^wt^ did not result in activation of AMPK in any cellular compartment (Fig. 2C). These results show that the energy defect caused by mtDNA depletion in Polɣ^D1135A^ cells primarily activates AMPK that is already associated with mitochondria.

The dynamics of the ensuing mitochondrial dysfunction in our experimental time course are summarized in Fig. 2E: the first effects on RC protein levels are observed on day 3 of induction, when the activation of mitochondria-associated AMPK also becomes apparent. Cell proliferation declines by day 5, and by day 6 the cells exhibit full-blown mitochondrial dysfunction involving diminished levels of most RC proteins, decreased MMP, and an overall decline in cellular energy state.

### AMPK helps maintain basal MMP, and stimulation of AMPK activity rescues the MMP loss in Polɣ^D1135A^ cells

Based on the timeline in Fig. 2E, we focused our efforts on Polɣ^D1135A^ cells on day 4 of induction, reasoning that this was a timepoint when the cells were suffering from mild mitochondrial dysfunction that could estimate the situation in mild to moderate mitochondrial disease. We asked whether pharmacological modulation of AMPK activity could alleviate some elements of the mitochondrial dysfunction, and treated uninduced or induced Polɣ^D1135A^ cells with the specific AMPK activator A-769662 for 48 or 72 h, as indicated at the top of Fig. 3A. Treatment with A-769662 resulted in the expected stimulation of AMPK activity, as evidenced by the increased phosphorylation of a major AMPK substrate acetyl-coenzyme A carboxylase (ACC) on serine-79, a site specifically phosphorylated by AMPK (Davies, Sim, and Hardie 1990) (Fig. S2A-B). The phosphorylation status of AMPK itself did not change appreciably following A-769662 treatment (Fig. S2C). Notably, A-769662 treatment increased mitochondrial membrane potential in a time-dependent manner, with 72-h treatment increasing MMP to up to 140% of starting levels in Polɣ^D1135A^-expressing cells (Fig. 3A). The impact of A-769662 on MMP was more pronounced in cells with a low starting MMP (80% and 140% increase in uninduced and induced Polɣ^D1135A^ cells after 72 h, respectively), and was sufficient to restore the MMP of induced Polɣ^D1135A^ cells to levels corresponding to the uninduced cells already after 48 h of treatment. We conclude that A-769662-mediated AMPK activation is sufficient to correct a considerable drop in mitochondrial membrane potential, and that the sustained effect of the activator can be observed at least 24 h after removal of the drug (Fig. 3A; 48 h treatment).

**Figure 3.**
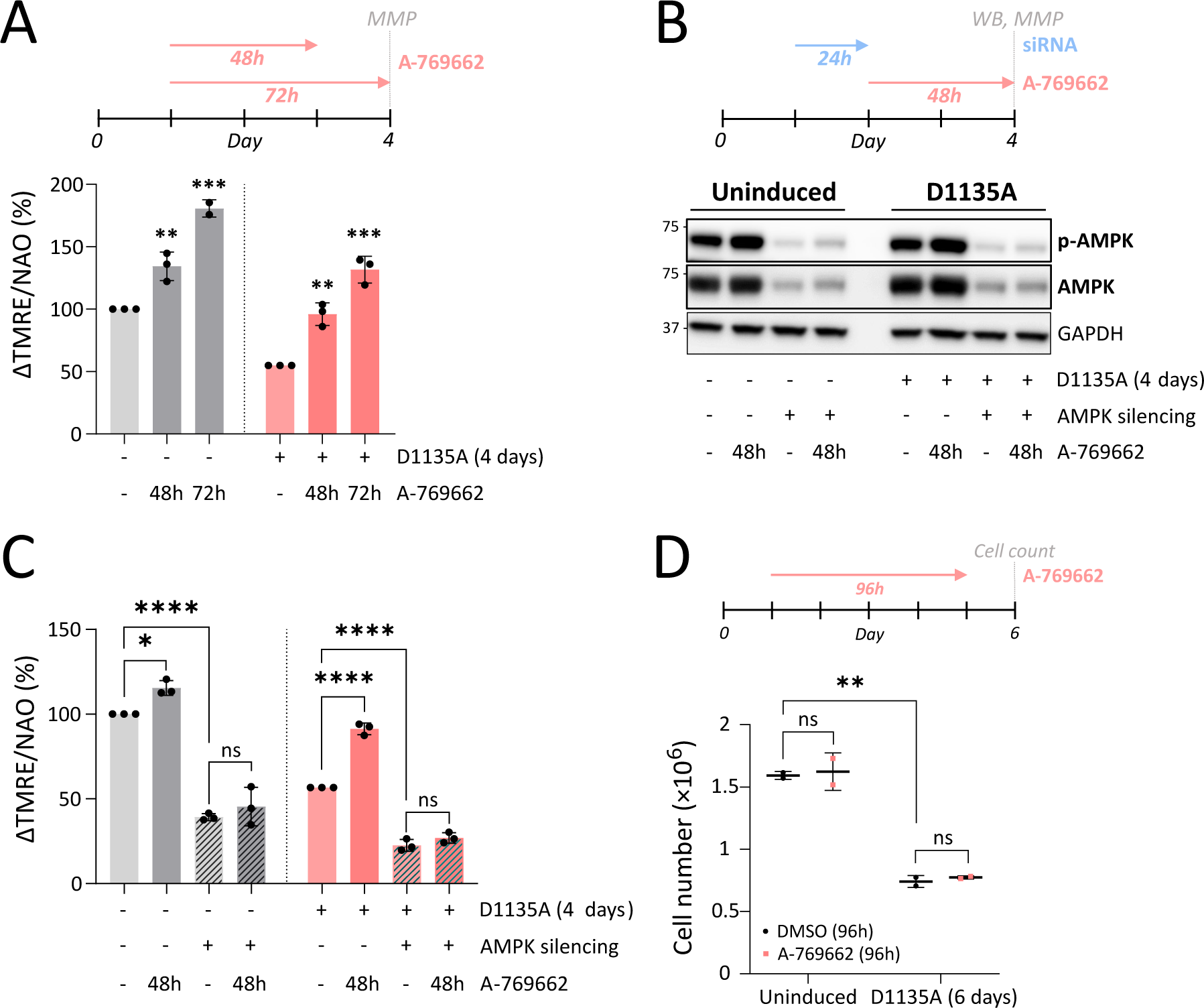
Pharmacological activation of AMPK restores MMP in mtDNA-depleted cells. **(A)** MMP in uninduced (*grey*) cells and cells expressing Polγ^D1135A^ (*pink*; 4-d induction) prior (-) and following AMPK activation with 100 µM A-769662 for 48 or 72 hours; n=3. **(B)** Immunoblotting indicates levels of phosphorylated AMPK (p-AMPK) and total AMPK in cells overexpressing PolG^wt^ and PolG^D1135A^ following silencing of AMPK using siRNA against α1/α2 AMPK. A representative image of n=3 is shown. **(C)** MMP in uninduced (*grey*) cells and cells expressing Polγ^D1135A^ (*pink*; 4-d induction) following AMPK silencing (*striped bars*) and/or A-769662 treatment (*darker shade*) from the experiment in Fig. 3B; n=3. **(D)** Cell proliferation assessed by cell number after a 96 h-treatment with 100 µM A-769662 (*pink*) in uninduced cells or ones induced to express Polγ^D1135A^ for 6 d; n=2. (n, number of biological replicates; ns, not significant; * P≤0.05; ** P≤0.01; *** P≤0.001; **** P≤0.0001).

In contrast to AMP-mimetic AMPK activators such as AICAR, A-769662 has been reported to be specific to AMPK (Xiao et al. 2013; Göransson et al. 2007). To confirm that the effect of A-769662 on MMP was AMPK-dependent, we next silenced the catalytic α-subunit of AMPK by siRNA treatment, and followed the impact of A-769662 on MMP in uninduced or induced Polɣ^D1135A^ cells. The siRNA treatment resulted in efficient knockdown of AMPKα as well as a clear drop in MMP in both uninduced and induced cells, indicating that AMPK is required for maintaining normal mitochondrial membrane potential (Fig. 3B-C). This finding is in good agreement with the reported impairment of mitochondrial respiration in mouse models lacking α1 and/or α2 subunits of AMPK (Lantier et al. 2014; Viollet et al. 2009). Further, as was seen in Fig. 3A, treatment with A-769662 increased the MMP in both the uninduced and induced cells, and the increase was more substantial in the induced cells that started from a lower MMP (Fig. 3C). However, A-769662 had no impact on the MMP in cells transfected with an siRNA against AMPK, confirming that the stimulatory effect of A-769662 on MMP is dependent on the presence of AMPK.

We have previously shown that the cell proliferation defect of *Saccharomyces cerevisiae* cells suffering from mitochondrial dysfunction can be rescued by increasing their MMP (Gorospe et al. 2022), and therefore now analyzed cell proliferation following A-769662 treatment in uninduced cells and ones induced for 6 days. As shown in Fig. 3D, the rescue in MMP observed after A-769662 treatment of induced Polɣ^D1135A^ cells was insufficient to improve cell proliferation, indicating that their growth defect was due to other consequences of the mitochondrial dysfunction than solely MMP loss.

### The A-769662-mediated activation of AMPK improves mtDNA copy number and respiratory chain protein levels

Given the positive impact of A-769962 on the mitochondrial membrane potential of induced Polɣ^D1135A^ cells, we sought to understand the molecular basis of this outcome by exploring the effect of A-769662 on the levels of mtDNA and RC subunits. A 72-h treatment of cells with A-769662 caused a small but statistically significant increase in the mtDNA copy number of induced Polɣ^D1135A^ cells, increasing mtDNA from 11% to 20% of the levels observed in uninduced cells (Fig. 4A). Interestingly, A-769662 treatment did not alter mtDNA levels in uninduced cells. Analysis of RC protein levels in induced cells revealed a positive effect of A-769662 on the analyzed CI, CIII and CIV subunits, but no alteration in CII or CV subunits that were only mildly depleted relative to uninduced cells to begin with (Fig. 4B-G). As was observed with mtDNA copy number, activator treatment had a differential effect on induced and uninduced cells: the protein levels of the latter only increased in one case (CI), while they decreased in two cases (CIII and CIV) and were not affected for CII and CV. Taken together, these results suggest that specific AMPK stimulation can alleviate the effects of mtDNA depletion and that the downstream effects of A-769662-mediated AMPK activation can vary, likely depending on the energy metabolic context of the cell.

**Figure 4.**
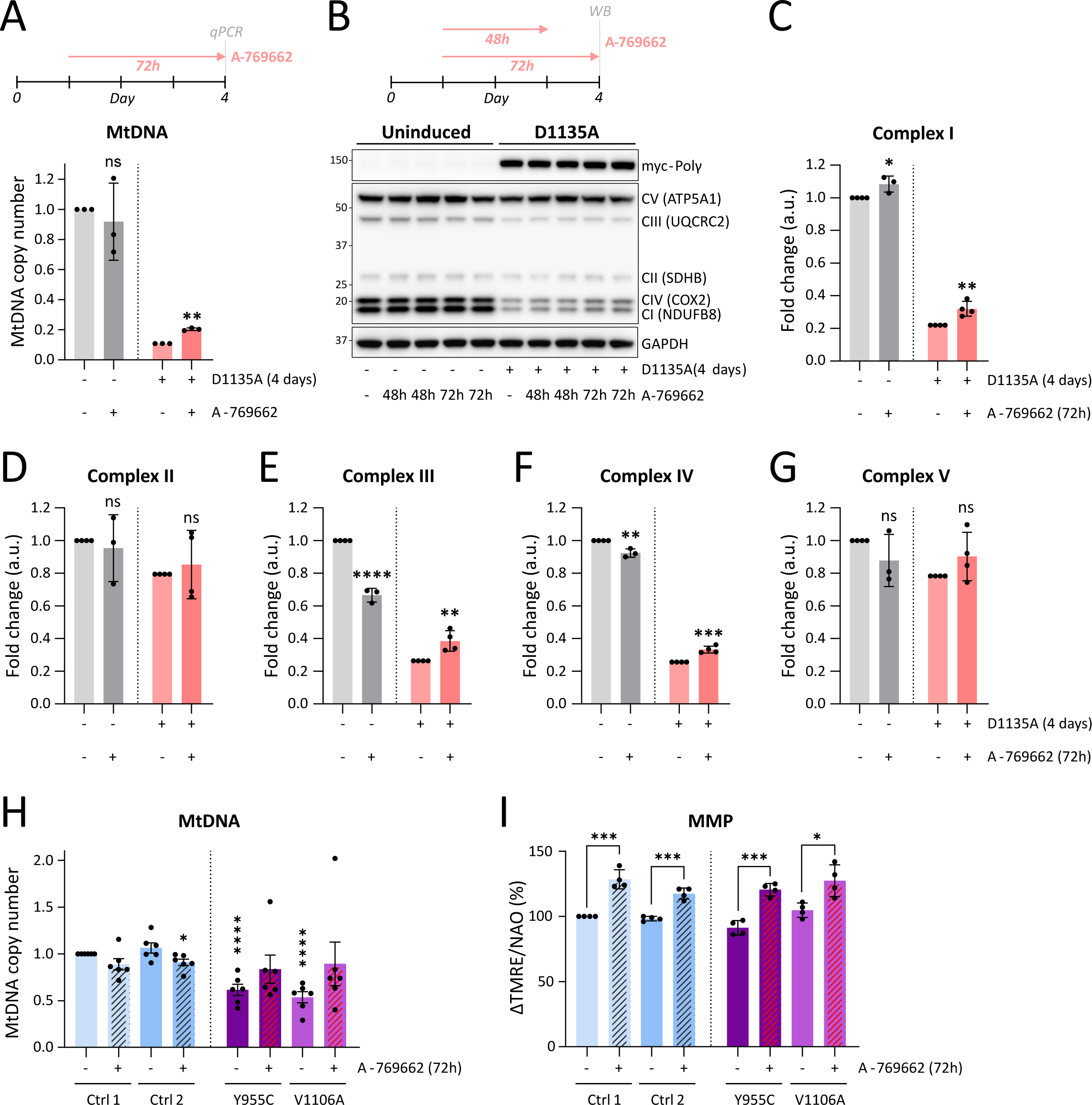
Pharmacological activation of AMPK ameliorates pathological phenotypes in cells expressing mutant variants of Polγ. **(A)** Relative mtDNA copy number after 72 h treatment with A-769662 (100 µM) reveals increased mtDNA levels in cells expressing Polγ^D1135A^ (*pink*) for 4 days compared to its respective DMSO control. n=3. **(B)** Expression of subunits of the mitochondrial respiratory complexes (CI-CV) after 48 h and 72 h treatment with A-769662 (100 µM) in uninduced cells and in cells expressing Polγ^D1135A^ (myc-Polγ) for 4 days. Protein levels were assessed by WB using an OXPHOS cocktail antibody. Protein molecular weight (kDa) are indicated on the left and protein names (and their respective respiratory complexes) are indicated on the right. A representative image of n=3-4 is shown. **(C, D, E, F and G)** Quantification of the blots shown in **(B)**; protein levels were normalized against GAPDH. **(H)** Relative mtDNA copy number from human primary fibroblast cells (2 controls; 2 patient cell lines bearing the Polγ homozygous mutations Y955C and V1106A, respectively) was measured before and after treatment with A-769662 (100 µM) for 24 h. n=6. Asterisks on untreated samples indicate the p-value of the comparison with the untreated Control 1 cell line; the asterisk on the A-769662-treated Ctrl2 sample marks the comparison with the untreated Ctrl2. **(I)** The mitochondrial membrane potential (MMP) of human primary fibroblast cells treated or not with A-769662 (100 µM) for 72 h was estimated by flow cytometry as described in Fig. 1K. Relative MMP is expressed as ΔTMRE/NAO. n=4. (n, number of biological replicates; ns, not significant; * P≤0.05; ** P≤0.01; *** P≤0.001; **** P≤0.0001).

So far, our experiments were carried out in cells expressing a dominant-negative Polɣ mutant with highly deleterious effects on mtDNA maintenance and mitochondrial function. Given that specific AMPK activation imparted some modest but promising effects even under this extreme scenario, we next applied A-769662 to cells with a milder and more clinically-relevant level of mtDNA depletion. To this end, we made use of fibroblasts derived from patients suffering from mitochondrial myopathy, as well as sex- and age-matched control cell lines. The mutant lines were homozygous either for PolɣA^V1106A^ or for the PolɣA^Y955C^ mutation that is the most common autosomal dominant mutation in *POLG* and a cause of progressive external ophthalmoplegia (PEO) (Rahman and Copeland 2019). Both mutant lines showed a depletion of mtDNA to approximately 60% of the levels found in control cells (Fig. 4H; 61% and 54% in Polɣ^Y955C^ and Polɣ^V1106A^ cells, respectively). Treatment with A-769662 led to a slight increase that lacked statistical significance in patient cell lines, and no change or a decrease in the control lines (Fig. 4H). However, A-769662-treatment resulted in an elevation of MMP, indicating that it enhanced mitochondrial function (Fig. 4I). Unlike the observations in uninduced *vs.* induced Polɣ^D1135A^ cells in Fig. 3A, the impact of A-769662 was comparable in control and patient cell lines, with treatment on average increasing MMP to 125% of starting values.

Taken together, the experimental findings in Figures 3-4 indicate that specific AMPK activation using A-769662 has a beneficial impact on mitochondrial function not only in the Flp-In T-REx 293 model system with severe mtDNA depletion, but also in mitochondrial myopathy-associated cells (Fig. 3A; Fig. 4I). However, the mechanisms mediating the positive effects of pharmaceutical AMPK activation differed between our experimental models: while the increased MMP in the Polɣ^D1135A^-expressing cells was at least partially mediated by an increase in mtDNA copy number (Fig. 4A), A-769662 did not significantly improve mtDNA levels in patient fibroblasts with less severe mtDNA depletion (Fig. 4H). The results of this study warrant expanding the repertoire of potential therapeutic applications of specific AMPK activators to include mitochondrial defects caused by mtDNA depletion.

## Discussion

Given its role as a master metabolic regulator tasked with ensuring energy homeostasis and adaptability to changing metabolic demands, AMPK naturally serves as a guardian of mitochondrial integrity and function. This multifaceted role involves regulation of mitochondrial biogenesis, dynamics and morphology as well as the quality control of mitochondria through mitophagy (Herzig and Shaw 2018). AMPK signaling also helps determine mitochondrial mobility in neuronal axons, essentially concentrating mitochondria at sites of energetic stress (Watters et al. 2020).

In the current study, we concretely show that AMPK activity is essential for maintaining MMP in healthy cells, and that mitochondria-associated AMPK is activated early on during the progression of mitochondrial dysfunction, in line with its established role in restoring energy homeostasis (Fig. 2; Fig. 3C). Here it is relevant to note that the phosphorylation level of cytosolic AMPK was not affected, and that assessment of total cellular AMPK would yield a very different result in terms of the timing of activation than that on mitochondria-associated AMPK (AMPK activation on day 9 *vs.* day 3 in total extracts and mitochondrial fractions, respectively). Further promotion of AMPK activity using the specific small-molecule activator A-769662 was sufficient to recover normal MMP in cells suffering from severe mitochondrial dysfunction following mtDNA depletion (Fig. 3). Although A-769662 treatment improved many features of the induced Polɣ^D1135A^ cells including mtDNA copy number and RC protein levels (Fig. 3-4), it did not correct the proliferation defect of these cells (Fig. 3D). In light of past work connecting AMPK activation to a G1/S checkpoint through the p53-p21-cyclin E axis (Mitra et al. 2009; Jones et al. 2005; Mandal et al. 2005), the persisting proliferation defect in AMPK-active cells is not entirely surprising. On the other hand, the proliferation defect was not solely maintained by AMPK, since inhibiting AMPK alone with the small-molecule inhibitor dorsomorphin was insufficient to rescue proliferation (Fig. S2D-H). It remains to be seen whether simultaneous inactivation of AMPK and the correction of MMP through an AMPK-independent mechanism could be a feasible approach to restore normal proliferation in cells with severe mtDNA depletion.

To our knowledge, this is the first report of a stimulatory effect of A-769662 on mtDNA copy number. Previous findings regarding the effects of AMPK agonists on mtDNA levels have come to disparate results. While AMPK activation via administration of the AMP-mimetic activator AICAR did not impact mtDNA levels in mice with RC defects (Peralta et al. 2016; Viscomi et al. 2011), treatment of HEK293T cells with compound 991 increased copy number by ∼ 1.3-fold (Malik et al. 2023). It should however be noted that all these three studies were carried out in models with a normal starting level of mtDNA, while in our study the largest effects of A-769662 on copy number were seen in the induced Polɣ^D1135A^ cells with extreme mtDNA depletion (ca 15% mtDNA remaining), while control cells or patient fibroblasts with milder depletion (ca 60% mtDNA remaining) showed no increase with A-769662 treatment (Fig. 4A and H). The actual impact of the various AMPK activators on mtDNA copy number can therefore not be directly compared across the different studies, and should be more thoroughly investigated in the future. Regardless, our findings call for exploration of the utility of specific AMPK activators as a potential experimental and/or therapeutic avenue even in disorders caused by mtDNA depletion.

The observed differential effects of A-769662 on control cells *vs.* Polɣ^D1135A^-expressing cells with mtDNA depletion are intriguing. Why does AMPK activation lead to different outcomes depending on the cellular context? One possible explanation is that in mtDNA depleted cells, the increased AMP or ADP levels contribute to co-activation of AMPK such that a higher activity is reached than with A-769662 alone. This hypothesis is supported by the fact that AMP and A-769662 bind different sites of the heterotrimeric AMPK complex — while AMP binds the cystathionine-β-synthase (CBS) domains on the regulatory ɣ-subunit, A-769662 binds the allosteric drug and metabolite-binding site (ADaM) located between the kinase domain of the catalytic α-subunit and the carbohydrate-binding module of the β-subunit (Xiao et al. 2013). Bultot et al. observed considerable co-activation of AMPK complexes derived from mouse hepatocytes or C_2_C_12_ myotubes using activators that bound to these two distinct sites (Bultot et al. 2016), so we speculate that the same may be occurring in the Polɣ^D1135A^-expressing cells treated with A-769662 in our study. The fact that no preferential MMP increases were seen in the patient-derived fibroblasts over controls in Fig. 4I is well in line with this reasoning. Although we did not measure the adenine nucleotide pools in the patient fibroblasts, their MMP is indistinguishable from that of the control cells, suggesting that their mitochondrial ATP production and thus cellular energy status is in the normal range.

In humans, each of the three AMPK subunits is present in multiple isoforms (two isoforms each of α and β, three of ɣ), theoretically giving rise to up to 12 distinct heterotrimeric AMPK complexes. The different AMPK complexes may vary in terms of tissue-specificity, subcellular localization and/or response to stress stimuli (Herzig and Shaw 2018). A shortcoming of A-769662 in the context of targeting mitochondrial stress is that it only activates AMPK complexes containing a β1-, but not a β2-subunit (Scott et al. 2014), and is therefore not expected to exert an effect on AMPK complexes associated with skeletal muscle mitochondria that primarily contain β2-subunits (Drake et al. 2021). In contrast, compound 991 (also known as ex229) binds the same ADaM site on AMPK as A-769662 but is 5-10 times more potent and shows no isoform specificity regarding the β-subunit (Lai et al. 2014; Xiao et al. 2013), highlighting it as the preferential AMPK agonist for further studies.

The targeting of a central regulator like AMPK as a treatment strategy comes with its own risks and challenges. In some studies, the chronic activation of AMPK has shown undesirable side effects such as cardiac and kidney hypertrophy and Alzheimer’s disease (Wilson et al. 2021; Zimmermann et al. 2020). However, clinical trials and animal studies with other AMPK agonists did not reveal harmful side effects, indicating that activating this central regulator is feasible and safe with the right compounds (López-Pérez et al. 2021; Pinkosky et al. 2016; Ray et al. 2024; Cusi et al. 2021). In conclusion, the positive effects of AMPK stimulation on mtDNA levels and/or mitochondrial membrane potential in cell lines with mild to severe mtDNA depletion broaden the potential applications of small-molecule AMPK agonists, and support further exploration of the effects of this group of compounds in the context of mitochondrial dysfunction.

## Acknowledgements

We thank Dr. Johannes Spelbrink (Radboud University Medical Center, Nijmegen, The Netherlands) for the Flip-In T-REx 293 cells, and the Protein Expertise Platform at Umeå University (a part of Protein Production Sweden) for cloning assistance. This work was supported by grants from the Swedish Research Council (grant number 2019-01874), The Swedish Cancer Society (19 0022 JIA, 22 2381 Pj), The Knut and Alice Wallenberg Foundation (KAW 2021.0053), The Swedish Society for Medical Research (S17-0023) and the Åke Wiberg Foundation (M20-0132) to P.H.W. R.D.S.P. is funded by The Lily Foundation, Muscular Dystrophy UK (MDUK), a seedcorn award from the Rosetrees Trust and Stoneygate Foundation and Medical Research Council (UK) strategic award MR/S005021/1 to establish an International Centre for Genomic Medicine in Neuromuscular Diseases (ICGNMD). R.D.S.P. and M.F. are supported by a Medical Research Council (UK) Clinician Scientist Fellowship (MR/S002065/1), Medical Research Council (UK) award MC_PC_21046 to establish a National Mouse Genetics Network Mitochondria Cluster (MitoCluster), and the LifeArc Centre to Treat Mitochondrial Diseases (LAC-TreatMito). The funding sources were not involved in study design, data collection or analysis, or in the decision to publish.

## Author contributions

GC-conceptualization, investigation, methodology, formal analysis, data curation, validation, visualization, project administration; BR, TVHN, JMEF, FR, ICM-investigation, methodology, formal analysis, validation; MF, RDSP-conceptualization, resources; MD-investigation, validation; SW-methodology, supervision, funding acquisition; AH – methodology, formal analysis, supervision; PHW-conceptualization, formal analysis, data curation, funding acquisition, project administration, supervision, writing-original draft. All authors reviewed and edited the final version.

