## Supplemental figures 1-2 for "The specific AMPK activator A-769662 ameliorates pathological phenotypes following mitochondrial DNA depletion"

**Supplemental figure legends**

**Supplemental Figure 1, connected to Figures 1-2. Effects of the progressive mitochondrial dysfunction induced by Polγ$^{D1135A}$ expression. (A, B)** Expression of Polγ$^{wt}$ and Polγ$^{D1135A}$ (myc-Polγ) after induction by doxycycline in a dose dependent manner (3 and 20 ng/mL). No leaky expression is observed in the absence of doxycycline; n=1. **(C)** MtDNA copy number is undetectable in HEK293 T-Rex $\rho^0$ cells compared to $\rho^+$ cells; n=2. **(D)** Mitochondrial membrane potential in $\rho^+$ and $\rho^0$ cells was assessed by flow cytometry and is expressed as ΔTMRE/NAO; n=2. **(E, F)** Levels of dNTPs and rNTPs are shown for uninduced cells (*0 d*) and cells expressing Polγ$^{wt}$ (*grey*) or Polγ$^{D1135A}$ (*pink*) for 3 or 6 days (DOI = days of induction); n=4. **(G)** Immunoblotting shows levels of phosphorylated ATM (p-ATM) and total ATM in cells expressing Polγ$^{WT}$ and Polγ$^{D1135A}$ for 3, 6 and 9 days, and in $\rho^0$ cells as well as in control cells treated with UV-light. Quantification of ATM phosphorylation is shown in the lower panel; n=2. **(H, I)** Cell cycle phases (G1, S and G2) were assessed by flow cytometry in cells expressing Polγ$^{wt}$ and Polγ$^{D1135A}$ for 3, 6 and 9 days; n=2. Cells treated with the mitochondrial uncoupler CCCP for 16h are shown as a control and show a similar increase in S phase cells; n=1. **(J)** Cell death was measured in cells expressing Polγ$^{wt}$ and Polγ$^{D1135A}$ for 6 days; n=2. Treatment with camptothecin (CPT), a topoisomerase inhibitor that induces DNA damage, was included as a positive control for cell death; n=1. (n, number of biological replicates; * $P \leq 0.05$; ** $P \leq 0.01$; *** $P \leq 0.001$; **** $P \leq 0.0001$)

**Supplemental Figure 2, connected to Figures 3-4. Additional effects of the pharmaceutic modulation of AMPK activity. (A)** Western blots show the levels of p-ACC, ACC, p-AMPK and AMPK in cells treated with 200 μM A-769662 for the indicated time period. β-Tubulin was used as loading control. (n=1) **(B, C)** Quantification of blots from Fig. S2A confirm the induction of AMPK activity, as judged by the p-ACC/ACC ratio, following A-769662 treatment. The treatment caused no appreciable change in the pAMP/AMPK ratio. **(D)** Western blots showing the levels of p-ACC, ACC, p-AMPK and AMPK in cells treated with 20 μM of the AMPK inhibitor dorsomorphin for the indicated time periods. β-Tubulin was used as loading control; (n=1). **(E, F)** Quantification of blots in Fig. S2D shows that dorsomorphin treatment decreased AMPK activity as judged by the ratio of pACC to ACC, but caused no appreciable change to the AMPK phosphorylation status (pAMPK/AMPK). **(G)** AMPK inhibition by dorsomorphin (20 μM) alters MMP in uninduced (*grey*) cells and in cells expressing Polγ$^{D1135A}$ (*blue*). (n=3) **(H)** AMPK inhibition impairs normal cell proliferation and exacerbates the effect of Polγ$^{D1135A}$ after 6 days of induction with doxycycline. n=2. (n, number of biological replicates; * $P \leq 0.05$; ** $P \leq 0.01$; *** $P \leq 0.001$; **** $P \leq 0.0001$).

### Supp 1

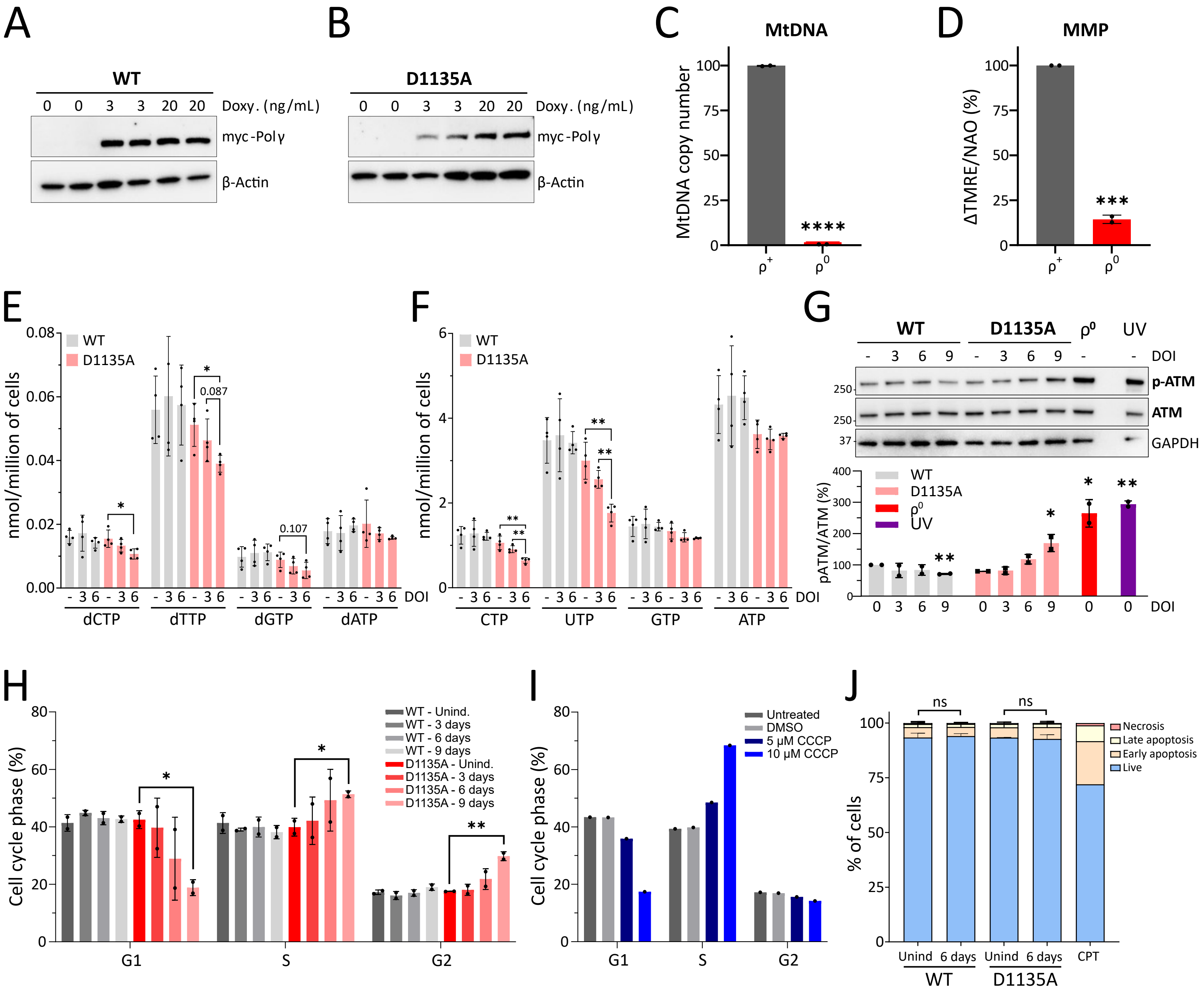

A
WT
0 0 3 3 20 20 Doxy. (ng/mL)
myc-Polγ
β-Actin
B
D1135A
0 0 3 3 20 20 Doxy. (ng/mL)
myc-Polγ
β-Actin
C
MtDNA
MtDNA copy number
****
ρ+ ρ0
D
MMP
ΔTMRE/NAO (%)
***
ρ+ ρ0
E
WT
D1135A
nmol/million of cells
0.087
0.107
DOI
dCTP dTTP dGTP dATP
F
WT
D1135A
nmol/million of cells
DOI
CTP UTP GTP ATP
G
WT D1135A ρ0 UV
DOI
p-ATM
ATM
GAPDH
pATM/ATM (%)
WT
D1135A
ρ0
UV
H
Cell cycle phase (%)
WT - Unind.
WT - 3 days
WT - 6 days
WT - 9 days
D1135A - Unind.
D1135A - 3 days
D1135A - 6 days
D1135A - 9 days
G1 S G2
I
Cell cycle phase (%)
Untreated
DMSO
5 μM CCCP
10 μM CCCP
G1 S G2
J
% of cells
ns
ns
Necrosis
Late apoptosis
Early apoptosis
Live
Unind 6 days Unind 6 days CPT
WT D1135A

### Supp 2

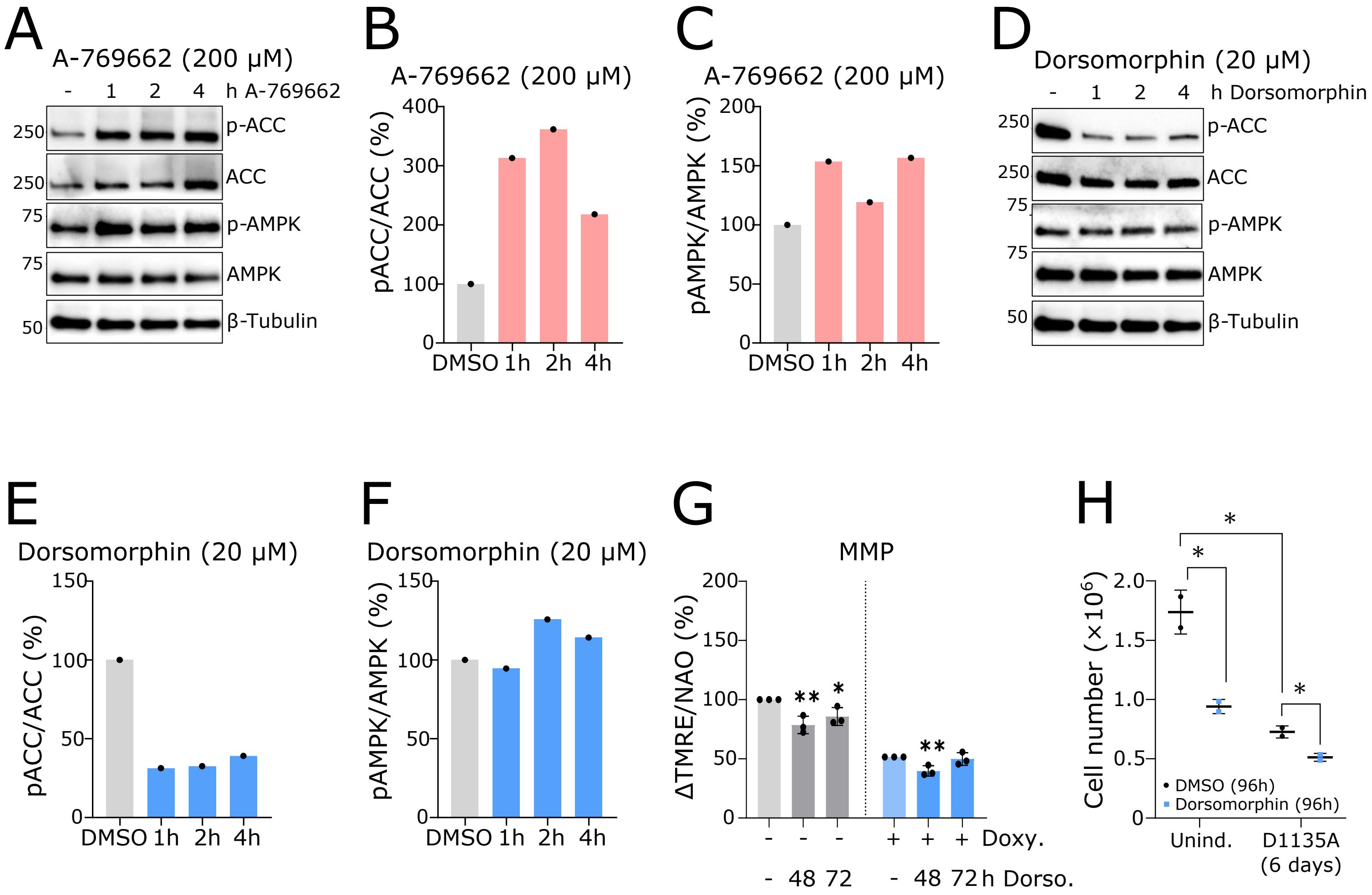

A
A-769662 (200 µM)
- 1 2 4 h A-769662
250 p-ACC
250 ACC
75 p-AMPK
75 AMPK
50 β-Tubulin
B
A-769662 (200 µM)
pACC/ACC (%)
DMSO 1h 2h 4h
C
A-769662 (200 µM)
pAMPK/AMPK (%)
DMSO 1h 2h 4h
D
Dorsomorphin (20 µM)
- 1 2 4 h Dorsomorphin
250 p-ACC
250 ACC
75 p-AMPK
75 AMPK
50 β-Tubulin
E
Dorsomorphin (20 µM)
pACC/ACC (%)
DMSO 1h 2h 4h
F
Dorsomorphin (20 µM)
pAMPK/AMPK (%)
DMSO 1h 2h 4h
G
MMP
ΔTMRE/NAO (%)
- - - + + + Doxy.
- 48 72 - 48 72 h Dorso.
H
Cell number (×10⁶)
DMSO (96h)
Dorsomorphin (96h)
Unind.
D1135A (6 days)
